# Multiple modes of DNA compaction by protamine

**DOI:** 10.1101/2023.12.08.570784

**Authors:** Vikhyaat Ahlawat, Huan-Xiang Zhou

## Abstract

In sperm cells, protamine replaces histones to compact DNA 10-20 times more than in somatic cells. To characterize the extreme compaction, we employed confocal microscopy and optical tweezers to determine the conformations and stability of protamine-bound λ-DNA. Confocal images show increasing compaction of λ-DNA at increasing protamine concentration. In the presence of protamine, single λ-DNA molecules form bends and loops that unravel at 10-40 pN forces as well as coils that shorten the contour length by up to 40% and withstand forces strong enough (∼55 pN) for strand separation. Strand separation nucleates coils, indicating protamine insertion into DNA bases. Protamine may participate in both local and higher-order chromatin organization, leading to extreme compaction and global transcription silencing.

**One-Sentence Summary:** Protamine compacts sperm DNA in multiple modes, producing bends and loops but also coils that may block transcription.

---

The packing of DNA inside a cell is critical to the regulation of gene expression (*1*). In somatic cells, compaction is a hierarchal process (*2*), where DNA is first organized into an array of nucleosomes, which consist of 147 base pairs of DNA wrapped around octamers of core histones. Linker histones then bind to core and linker DNA and promote nucleosome condensation (*3-5*). Chromatin loops can form when more distant loci are brought together, via either bridging or phase separation by chromatin-associated proteins (*2*). The next two higher-order chromatin units are topologically associating domains (TADs) and compartments. The former arises from active extrusion of DNA loops through the ring-shaped cohesion complex. The latter may arise through phase separation by, e.g., heterochromatin protein 1 (HP1) (*6, 7*), and separates active regions (“A” compartments) from inactive regions (“B” compartments) of the genome. Single-molecule techniques including optical tweezers (OT) and fluorescence imaging have been extensively used to study the roles of chromosomal proteins in the 3-dimensional (3D) organization of chromatin *(8)*. Multistep nucleosome unwrapping from core histones (*9-11*), stabilization of mononucleosomes by linker histones (*12*), and bridging of distal DNA loci by non-histone proteins including Alba (*13*), HMO1 (*14*), synaptonemal complex protein 3 (SYCP3) (*15*), cohesion (*16, 17*), and HP1α(*18*) have been reported.

In mature sperm cells, >90% of histones are replaced with protamines (*19*). These small (30-50 residues), basic, intrinsically disordered proteins compact genomic DNA 10-to 20-fold more than in somatic cells (*20*). The extreme compaction protects sperm DNA from damage and facilitates its delivery to the oocyte cytoplasm. A variety of imaging techniques have been used to investigate the shapes of DNA when compacted by protamine, including transmission electron (TEM) (*21*), atomic force (AFM) (*21-24*), and fluorescence (*25, 26*) microscopy. Based on TEM and AFM imaging, Hud et al. (*21*) proposed that protamine condenses sperm DNA into toroids containing up to 60 kilo-base pairs (kb). More recent AFM studies on shorter DNA have revealed a variety of other shapes, including single loops (*23*), multiple loops emanating from a central position, and stacks of loops (*24*). Single-molecule fluorescence imaging has shown DNA can be compacted into a near-zero length when fully covered by protamine (*25*). According to high-throughput chromosome conformation capture data, the 3D chromatin organization of mouse sperm is very similar to that of somatic cells, including the formation of TADs and compartments (*19, 27*). It is unclear whether protamine acts only locally like histones, or also plays roles in higher-order organization of sperm chromatin. In addition, transcription is silenced in sperm but the mechanism is still a matter of speculation.

Here we used confocal microscopy and optical tweezers to determine the conformations and stability of λ-DNA (48.5 kb) when compacted by salmon protamine (amino-acid sequence: M_1_PRRRRSSSR P_10_VRRRRRPRV S_20_RRRRRRGGR R_30_RR). The results demonstrate three distinct modes of DNA compaction, which explain the extreme level of compaction in sperm chromatin and suggest a possible mechanism for transcription silencing.

## Protamine can achieve 100% compaction of λ-DNA

As a simple demonstration of DNA compaction by protamine (*26*), we acquired confocal images of λ-DNA (2 ng/μL or 3 μM base pairs) stained with DRAQ5 (5 μM) and settled on glass slides, at 0, 0.5, 1, 2, 5, and 10 μM protamine. Free λ-DNA adopts extended conformations (Fig. 1A), with maximum dimensions (*D*_max_) of 14.1 ± 3.2 μm (mean ± SD; *n* = 16). At 0.5 μM protamine, λ-DNA condenses into a globular shape (Fig. 1B), with *D*_max_ at 3.5 ± 1.9 μm (*n* = 29). With increasing protamine concentrations, λ-DNA foci become more and more compact (Fig. 1C-F), down to *D*_max_ = 0.9 ± 0.1 μm (*n* = 26) at 10 μM protamine. The latter *D*_max_ is close to the width of the point spread point function of the dye; therefore 10 μM protamine appears to achieve nearly 100% compaction of λ-DNA. By doping protamine with a FITC-labeled variant, we verified that protamine is concentrated in λ-DNA foci (Fig. 1F).

**Fig. 1.**
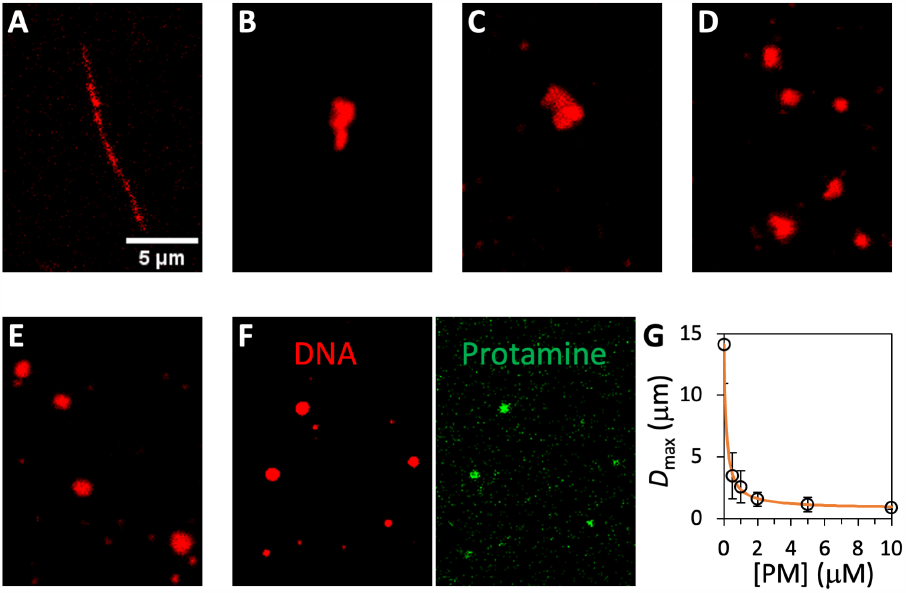
Confocal images showing λ-DNA condensation by protamine. (**A**) λ-DNA (3 μM base pairs) was stained with DRAQ5 (5 μM) settled on glass slides before confocal scanning. (**B**-**F**) Corresponding images at 0.5, 1, 2, 5, and 10 μM protamine, respectively. For (F), protamine was doped with protamine-FITC. Images from the red channel (from DNA staining) and the green channel (from protamine-FITC) demonstrate colocalization of DNA and protamine foci. (**G**) Decrease in maximum dimension (*D*_max_) with increasing protamine concentration ([PM]). Symbols display imaging data; a curve displays the fit to Eq. (S1) of Material and Methods.

The decrease in *D*_max_ with increasing protamine concentration is displayed in Fig. 1G. The data can be fit to a model where an increase in protamine concentration results in more coverage of the DNA molecule, and *D*_max_ is the length of the still not covered portion. A similar model was used by Brewer et al. (*25*) to analyze single DNA molecule compaction kinetic data, based on 100% compaction when the DNA molecule is fully covered by protamine.

## Protamine compacts single λ-DNA molecules in multiple modes

We tethered the ends of single λ-DNA molecules to two optically trapped beads and determined in the buffer channel the maximum extension (17.0 ± 0.4 μm (*n* = 23), measured at force ∼40 pN) before strand separation for the naked form. We then transferred the tether assembly to the protamine channel and moved trap 1 relative to a fixed trap 2, starting from an initial extension of 7.5 μm, at a constant speed of 1 μm/s (Fig. 2A inset) through successive stretch-relax cycles, separated by a waiting period, until the tethers broke. The force-extension curves revealed different modes of DNA compaction by protamine (Fig. 2 and fig. S1). In one mode (Fig. 2A), similar to what has been observed with other DNA-condensing proteins including core histone octamers (*9-11*), protamine appears to induce bends, which unravel under ∼15 pN of force. At high stretch, the rapid rise in the force-extension curve coincides with the counterpart for naked DNA, suggesting that the DNA stiffness is not affected by protamine in this compaction mode. Protamine stays bound after we brought the tether assembly back to the buffer channel, as the next stretch again shows the bending mode (fig. S2A).

**Fig. 2.**
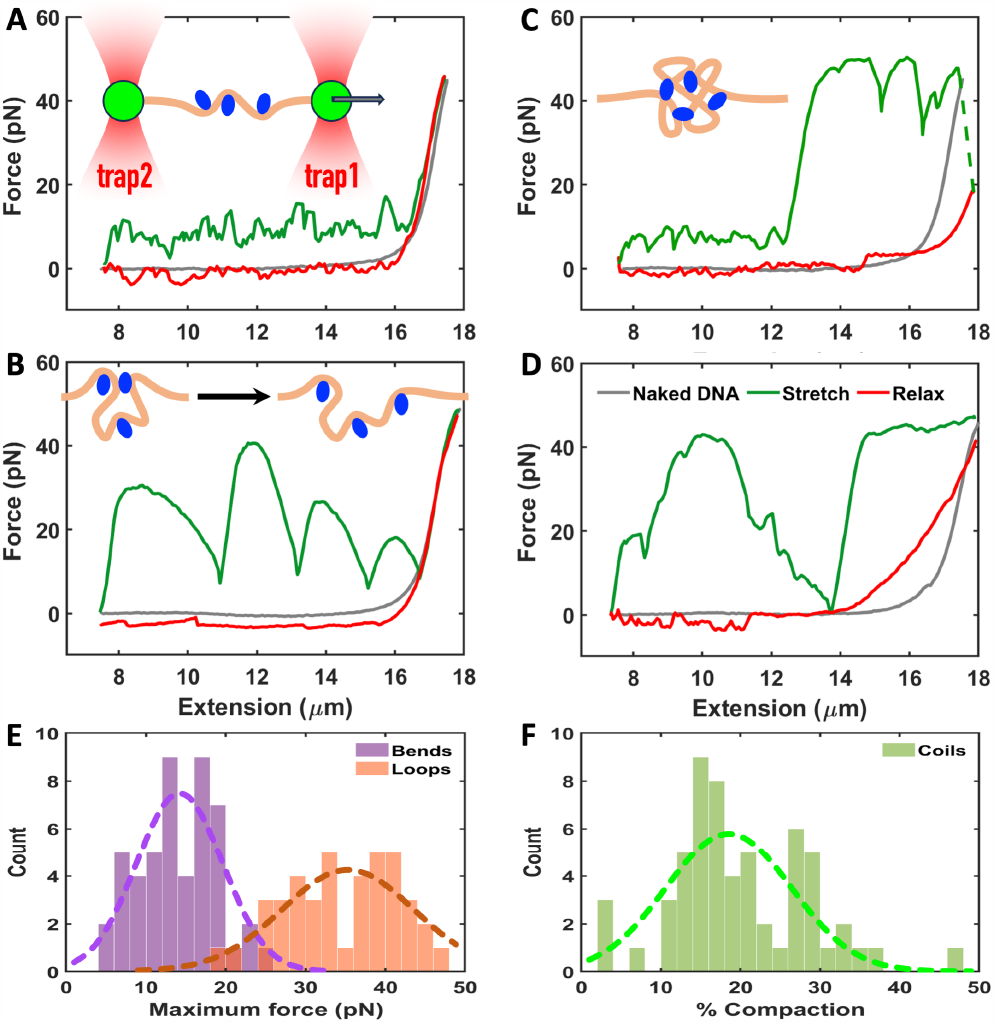
Force-extension curves demonstrating different modes of λ-DNA compaction by protamine. Protamine was at 10 μM. (**A**) Bending mode. Inset: illustration of a tethered DNA molecule with bends created by protamine molecules (blue ovals). (**B**) Looping mode. Inset: Breakup of a loop. (**C**) Coiling mode, with bends at low stretch. A large drop in force, indicated by a dashed line, occurs in the waiting period between the stretch and the relax. Inset: illustration of a coil. (**D**) Coiling mode, with loops at low stretch. (**E**) Histograms of maximum forces for breaking bends or loops. (**F**) Histogram for the percent reduction in DNA contour length by coils. In (E-F), Gaussian fits are shown as dashed curves.

In a second compaction mode, protamine appears to bridge two sites along the DNA molecule to create a loop (Fig. 2B inset). These loops unravel at forces up to 40 pN, each producing an increase of as much as ∼2 μm in extension (Fig. 2B). The bending mode and the looping mode can interconvert in successive stretch-relax cycles (fig. S3). After the loops are broken, the final rapid rise of the force-extension curve also coincides with the counterpart for naked DNA, again indicating a null effect of the bound protamine on DNA stiffness.

The third distinct mode of DNA compaction is signified by a much earlier rapid rise of the force-extension curve relative to naked DNA, followed by a plateau around 55 pN corresponding to strand separation (Fig. 2C). The shortened contour length, somewhat reminiscent of what was observed with HP1α (*18*) but to a much greater extent here, suggests that the DNA molecule forms coils that are stapled by tightly bound protamine molecules (Fig. 2C inset), such that they remain intact in the plateau, i.e., overstretched region with strand separation. Outside the coils, DNA molecules also form bends (Fig. 2C, low stretch region) or loops (Fig. 2D, low stretch region). Once formed, coils persist in subsequent stretch-relax cycles most of the time, again indicating tight binding of protamine. Indeed, after we transferred an overstretched DNA molecule containing coils to the buffer channel and relaxed it there, the subsequent stretch exhibits the coiling mode as if it stayed in the protamine channel (fig. S2B). Occasionally, coils partially unravel during the overstretch or the subsequent waiting period, as indicated by a sudden drop in pulling force and a concomitant increase in extension (Fig. 2C), but reform during most of the time the relax. Very rarely, we observed the complete unraveling of coils, such that the DNA molecule exhibits the bending mode in the next stretch (fig. S4).

We analyzed ∼50 force-extension curves in each mode to obtain histograms of the maximum forces for breaking bends or loops (Fig. 3E) or the percentage of reduction in contour length by coils (Fig. 3F). The maximum breaking forces are 14 ± 5 pN (*n* = 49) for bends and 35 ± 7 pN (*n* = 41) for loops; the percentages of compaction by coils are 20% ± 9% (*n* = 58), which are achieved at very high forces (> 40 pN). In comparison, *Xenopus* egg extracts, which presumably contain histones and all other condensing factors, produced a 60% compaction of a shorter (6.3 kb) DNA molecule at a minimal force (1 pN) (*28*). Both our confocal imaging (without force; Fig. 1) and Brewer et al.’s single-molecule imaging at a minimal force (*25*) suggest 100% DNA compaction by full protamine coverage.

**Fig. 3.**
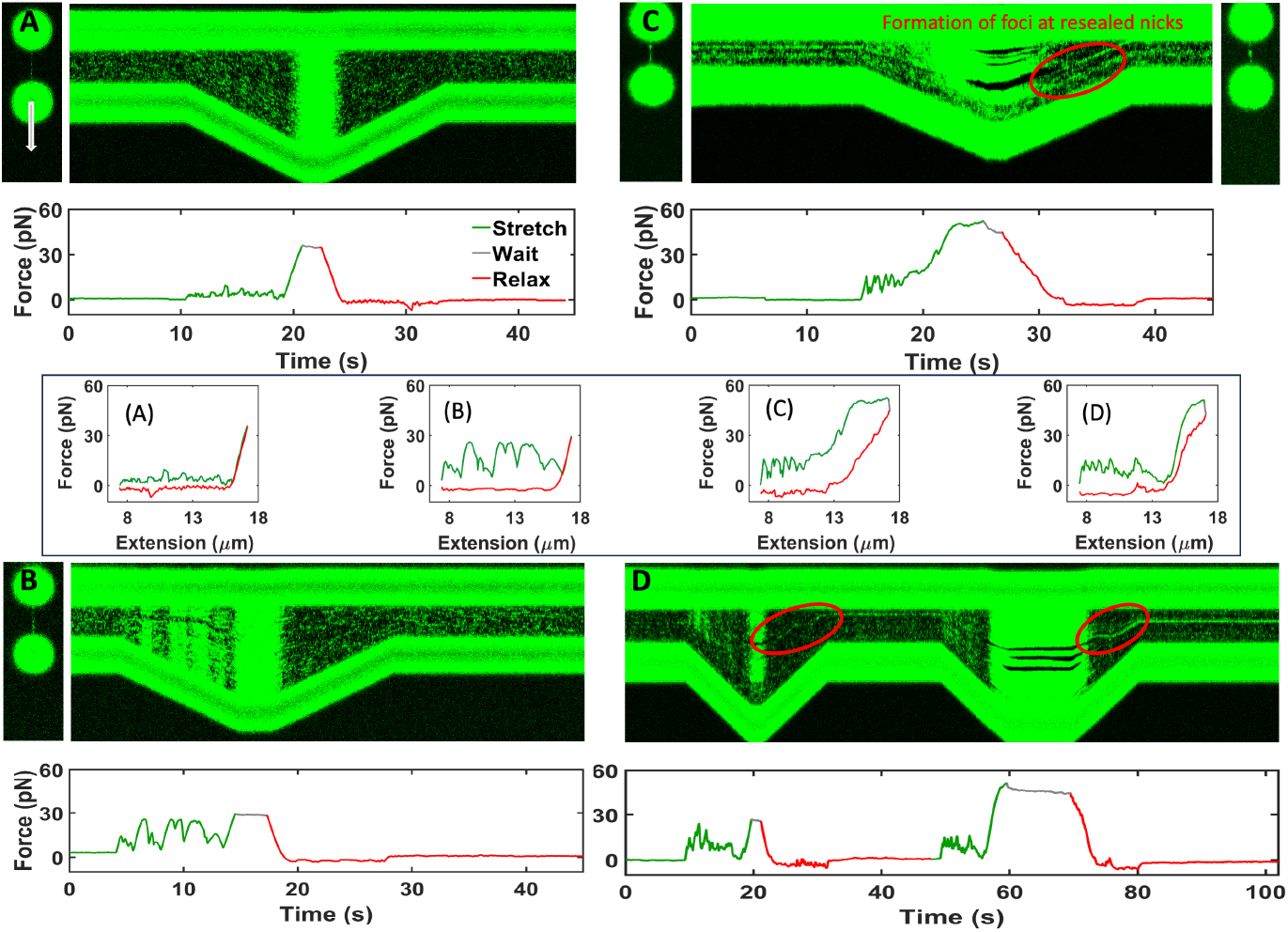
Kymographs illustrating the different compaction modes and an interconversion. SYTOX Orange (0.5 μM) was premixed in the protamine (10 μM) channel for DNA staining. (**A**) Top: the bending mode through a cycle from pre-stretch to stretch, waiting, relax, and back to pre-stretch; bottom: the corresponding trace of the pulling force. A 2D scan taken right before the kymograph is also shown. (**B**) Analogous results for the looping mode. (**C**) Analogous results for the coiling mode. SYTOX Orange foci are visible in both the kymograph and the pre- and post-kymograph 2D scans. (**D**) Conversion from the bending mode in one cycle to the coiling mode in the next. In (C) and (D), dark bands in the kymographs are due to nicks; red ovals highlight the emergence of SYTOX Orange foci at reannealed sites during the relax. Boxed middle row: force-extension curves corresponding to the force traces in the four panels; for panel (D), only the force-extension curve in the second cycle, i.e., the coiling mode, is shown.

## Strand separation nucleates coils

To gain molecular insight into the three compaction modes, we turned to correlative force-fluorescence measurements, by staining λ-DNA with SYTOX Orange (0.5 μM, present in the protamine channel), which exhibits very large fluorescence enhancement upon base intercalation (*29*). In the bending mode, SYTOX Orange binds to the DNA molecule uniformly at a low level in the pre-stretch period (during which the extension is held at 7.5 μm), and shows barely discernible increases in intercalation when forces are increased to straighten the bends but intense intercalation at high stretch (Fig. 3A). In the looping mode, there is a clear increase in intercalation each time when the force is increased to break a loop (Fig. 3B). By contrast, in the coiling mode, SYTOX Orange foci are already formed in the pre-stretch period, as shown by both a 2D scan and a kymograph (Fig. 3C); these foci most likely correspond to coils in which protamine is tightly bound. Moreover, dark bands emerge in the kymograph at high stretch, which unequivocally indicate nicks with strand separation (Fig. 3C). In the subsequent relax, the nicks reanneal, and SYTOX Orange foci then emerge from the reannealed sites, which signal that the coiling mode will be retained. Most interestingly, we captured on a kymograph the conversion from the bending mode in one stretch-relax cycle to the coiling mode in the next cycle (Fig. 3D). The conversion starts when nicks develop during the waiting period of the first stretch-relax cycle. Here again, when the nicks reanneal, SYTOX Orange foci emerge, signaling the coiling mode in the next cycle. This mode is indeed what we observed, where nicks develop early in the stretch and subsequently seed the formation of SYTOX Orange foci and DNA coils.

In fig. S5, we present another example of the conversion from the bending mode to the coiling mode. After the stretch in the bending mode, a kymograph captured first the development of nicks in the waiting period and then the emergence of SYTOX Orange foci during the relax (fig. S5A). The nicks and foci are also visible in 2D scans taken just before and after the kymograph, respectively. The subsequent stretch exhibits the coiling mode, as revealed by a time series of 2D scans (fig. S5B). The SYTOX Orange foci can be clearly identified in the first two frames. In the third frame, the dye intercalates the DNA molecule over its entire length. In the last three frames, nicks emerge along the DNA.

The data in Fig. 3C, D and fig. S5 show that strand separation generated at high stretch nucleates coils. To directly verify this conclusion, we wondered whether the bending mode could always be converted into the coiling mode by overstretching. We designed a seven-cycle sequence, with the first three to pick DNA molecules that are consistently in the bending (or looping) mode (Fig. 4A-C and fig. S6A-C). In the fourth cycle, the DNA is overstretched to introduce strand separation (Fig. 4D and fig. S6D). The last three cycles are then expected to consistently show the coiling mode. We tested this protocol seven times, and each time we indeed obtained the coiling mode in all the last three cycles (Fig. 4E-G and fig. S6E-G). Finally we even overstretched naked DNA in the buffer channel and then relaxed it in the protamine channel (Fig. 4H). The next stretch again produced the coiling mode.

**Fig. 4.**
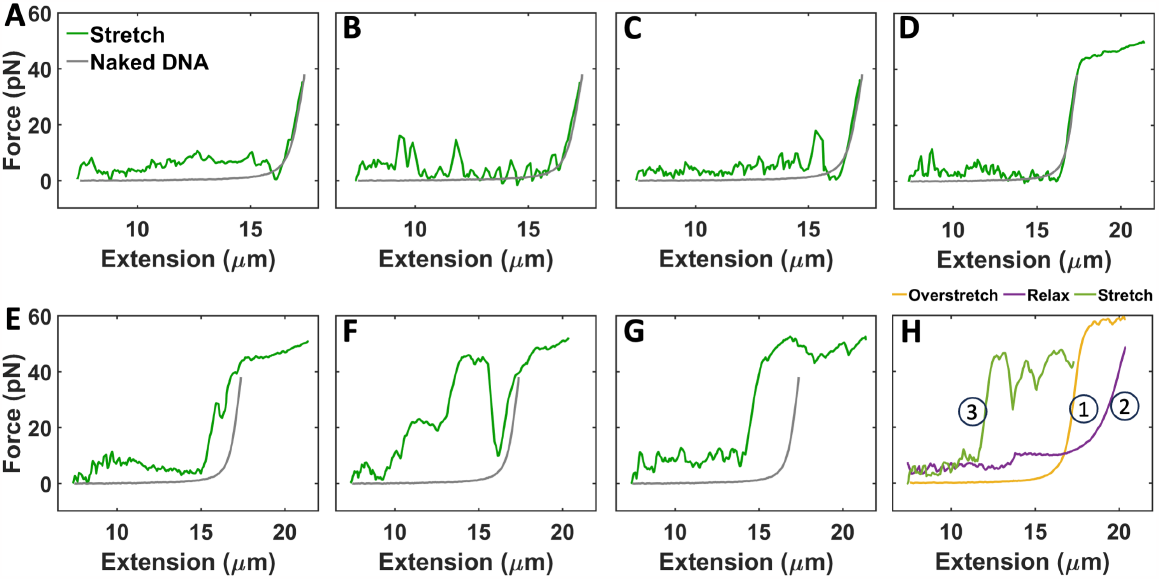
Induction of the coiling mode by overstretching DNA. (**A**-**G**) A seven-cycle sequence for inducing the conversion from the bending mode to the coiling mode. (A-C) present three successive regular stretch-relax cycles in the bending mode; overstretch is introduced in (D), resulting in the coiling mode in the subsequent cycles shown in (E-G). (**H**) Induction of the coiling mode by overstretching naked DNA.

## Coils may involve protamine-base interactions

Strand separation exposes DNA bases. The exposed bases allow for direct interaction with protamine, which may drive coil formation. Molecular dynamics simulations have shown that arginine sidechains of protamine form extensive interactions with molecular clusters of the nucleotide adenosine triphosphate (ATP), in the forms of salt bridges with their phosphate groups and cation-π interactions with their bases (*30*). These interactions drive the liquid-liquid phase separation of protamine-ATP mixtures. Support for direct protamine-base interaction in coils is provided by the effects of protamine on the unbinding and binding of SYTOX Orange. The fact that SYTOX Orange foci are found in the pre-stretch period of the coiling mode but not of the bending or looping mode (Fig. 3A-C) hints that protamine traps the intercalated dye in coils. To directly verify this conclusion, we overstretched DNA in the buffer channel and transferred it to the protamine channel (pre-mixed with SYTOX Orange) to induce the coiling mode. We then relaxed it to the pre-stretch state, where SYTOX Orange foci are present to confirm the coiling mode (fig. S7A, left panel). After transferring back to the buffer channel, the foci linger on the DNA molecule for > 35 s (fig. S7A, right panel). In contrast, after an overstretched naked DNA molecule with SYTOX Orange staining is transferred back to the buffer channel, the staining disappears immediately (fig. S7B).

We also wondered whether pre-bound protamine in coils could impede SYTOX Orange intercalation. To test this hypothesis, we again induced the coiling mode by transferring overstretched DNA from the buffer channel to the protamine channel, but this time without premixing SYTOX Orange. After confirming the coiling mode by the force-extension curve, we transferred this DNA molecule in the overstretched state to the dye channel, where it shows only intermittent staining (fig. S8A), in contrast to the complete, continuous staining of unnicked regions of DNA when the dye is premixed with protamine (overstretched regions in Fig. 3C, D and fig. S7A) or when an overstretched naked DNA molecule is transferred to the dye channel (fig. S8B).

Taken together, these observations show that protamine bound in coils acts as a barrier for an intercalating dye to get into and get off from the bases. Because the dye molecule is completely buried between two adjacent base pairs upon intercalation, it is most likely that protamine has to directly interact with the bases in order to interfere with dye intercalation.

## Protamine may participate in multiple levels of sperm chromatin organization

The single-molecule force spectroscopy and correlative force-fluorescence measurements have revealed that protamine compacts DNA in multiple modes. The bending mode, with breaking forces at ∼15 pN, is similar to the behavior exhibited by core histone octamers (*9-11*). The looping mode, with breaking forces at ∼40 pN, is analogous to the effects of Alba (*13*) and HMO1 (*14*). The coiling mode withstands forces strong enough (∼55 pN) for strand separation and substantially shortens the contour length, reminiscent of the results of SYCP3 (*15*) and HP1α (*18*). It is remarkable that a 33-residue disordered protein can compact DNA in such a multitude of ways and to such a high level.

We expect protamine to form distinct interactions with DNA in the different compaction modes. Because coils can withstand much higher forces than bends and loops, the protamine-DNA interaction in the former mode must be much stronger and likely involves more protamine molecules than in the latter two modes. A possible analogy for the distinct protamine-DNA interactions is provided by proteins that bind to DNA at a sequence-specific site with a very high affinity but also at nonspecific sites with moderate affinities (*31-34*). In particular, arginines in those proteins may insert more deeply into the DNA and form base-specific hydrogen bonds at the specific site, but form electrostatic interactions with the phosphate backbone at nonspecific sites. Although protamine is not known to have sequence specificity, in the coiling mode, its arginines may reach more deeply into the bases to form hydrogen bonds or even cation-π interactions (Fig. 5A). The observation that strand separation nucleates coils supports this notion; this aspect is reminiscent of the preference of linker histone H1 for DNA forks (*5*). Further evidence for protamine-base interaction is provided by the observation that protamine bound in coils impedes both the base intercalation and the release of SYTOX Orange.

**Fig. 5.**
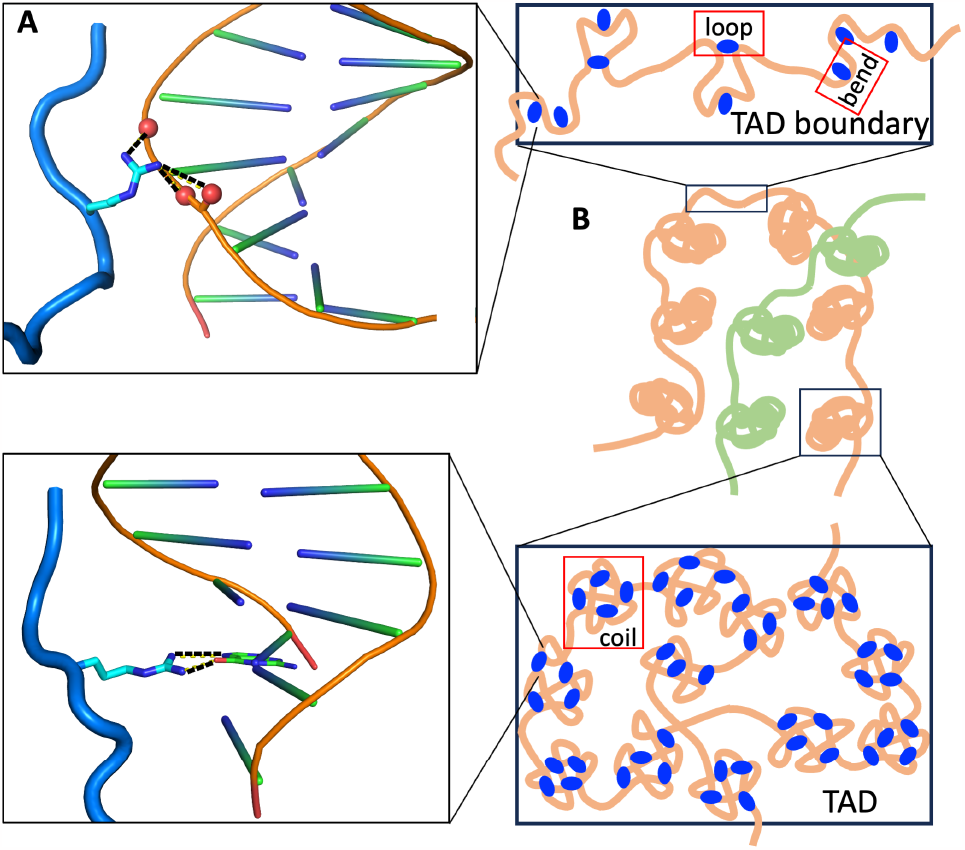
Protamine participates in multiple levels of sperm chromatin organization. (**A**) Protamine may interact only with backbone phosphates (top panel) in the bending and looping modes but also with bases (bottom panel) in the coiling mode. (**B**) Protamine compacts sperm chromatin locally by producing bends and at an intermediate level by producing loops. At an even higher level, protamine condenses TADs by producing coils.

The multiple modes of DNA compaction suggest that protamine participates not only in local but also higher-order organization of sperm chromatin. The effects of the bending mode correspond to those produced by histones. In addition, the looping mode may produce chromatin loops. The typical length of loops produced by protamine binding is ∼2 μm (Fig. 2B), corresponding to 5 kb of DNA. Most importantly, protamine binding produces coils that may populate in and condense TADs (Fig. 5B). The resulting extreme compaction can repress transcription, but protamine also provides an additional mechanism: coils are not breakable by forces exerted by RNA polymerases and will thus completely block transcription. This putative role explains why protamine is enriched in A compartments relative to B compartments in the mouse sperm chromatin (*19*).

## Acknowledgments

We thank Divya Kota for preparing protamine-FITC.

## Author contributions

Conceptualization: HXZ, VA

Methodology: VA

Investigation: VA, HXZ

Visualization: VA, HXZ

Funding acquisition: HXZ

Project administration: HXZ

Supervision: HXZ

Writing – original draft: HXZ, VA

Writing – review & editing: HXZ, VA

## Data and materials availability

All data are available in the main text or the supplementary materials, or from the authors upon reasonable request.

## References and Notes

1. A. Pombo, N. Dillon, Three-dimensional genome architecture: players and mechanisms. Nat Rev Mol Cell Biol 16, 245–257 (2015).

2. T. Misteli, The Self-Organizing Genome: Principles of Genome Architecture and Function. Cell 183, 28–45 (2020).

3. A. L. Turner et al., Highly disordered histone H1-DNA model complexes and their condensates. Proceedings of the National Academy of Sciences 115, 11964–11969 (2018).

4. B. A. Gibson et al., Organization of Chromatin by Intrinsic and Regulated Phase Separation. Cell 179, 470–484.e421 (2019).

5. R. Leicher et al., Single-stranded nucleic acid binding and coacervation by linker histone H1. Nat Struct Mol Biol 29, 463–471 (2022).

6. A. G. Larson et al., Liquid droplet formation by HP1α suggests a role for phase separation in heterochromatin. Nature 547, 236–240 (2017).

7. A. R. Strom et al., Phase separation drives heterochromatin domain formation. Nature 547, 241–245 (2017).

8. I. Heller, T. P. Hoekstra, G. A. King, E. J. Peterman, G. J. Wuite, Optical tweezers analysis of DNA-protein complexes. Chem Rev 114, 3087–3119 (2014).

9. B. D. Brower-Toland et al., Mechanical disruption of individual nucleosomes reveals a reversible multistage release of DNA. Proc Natl Acad Sci U S A 99, 1960–1965 (2002).

10. C. Diaz-Celis et al., Assignment of structural transitions during mechanical unwrapping of nucleosomes and their disassembly products. Proc Natl Acad Sci U S A 119, e2206513119 (2022).

11. L. H. Pope et al., Single chromatin fiber stretching reveals physically distinct populations of disassembly events. Biophys J 88, 3572–3583 (2005).

12. S. Rudnizky et al., Extended and dynamic linker histone-DNA Interactions control chromatosome compaction. Mol Cell 81, 3410–3421 (2021).

13. N. Laurens et al., Alba shapes the archaeal genome using a delicate balance of bridging and stiffening the DNA. Nat Commun 3, 1328 (2012).

14. D. Murugesapillai et al., DNA bridging and looping by HMO1 provides a mechanism for stabilizing nucleosome-free chromatin. Nucleic Acids Res 42, 8996–9004 (2014).

15. J. L. Syrjanen et al., Single-molecule observation of DNA compaction by meiotic protein SYCP3. eLife 6, e22582 (2017).

16. P. Gutierrez-Escribano et al., A conserved ATP- and Scc2/4-dependent activity for cohesin in tethering DNA molecules. Sci Adv 5, eaay6804 (2019).

17. Y. Kim, Z. Shi, H. Zhang, I. J. Finkelstein, H. Yu, Human cohesin compacts DNA by loop extrusion. Science 366, 1345–1349 (2019).

18. M. M. Keenen et al., HP1 proteins compact DNA into mechanically and positionally stable phase separated domains. eLife 10, e64563 (2021).

19. Y. H. Jung et al., Chromatin States in Mouse Sperm Correlate with Embryonic and Adult Regulatory Landscapes. Cell Rep 18, 1366–1382 (2017).

20. W. S. Ward, D. S. Coffey, DNA packaging and organization in mammalian spermatozoa: Comparison with somatic cells. Biol Reprod 44, 569–574 (1991).

21. N. V. Hud, M. J. Allen, K. H. Downing, J. Lee, R. Balhorn, Identification of the Elemental Packing Unit of DNA in Mammalian Sperm Cells by Atomic Force Microscopy. Biochem Biophys Res Commun 193, 1347–1354 (1993).

22. M. J. Allen, E. M. Bradbury, R. Balhorn, AFM analysis of DNA-protamine complexes bound to mica. Nucleic Acids Res 25, 2221–2226 (1997).

23. O. A. Ukogu et al., Protamine loops DNA in multiple steps. Nucleic Acids Res 48, 6108–6119 (2020).

24. R. B. McMillan et al., Protamine folds DNA into flowers and loop stacks. Biophys J 122, 4288–4302 (2023).

25. L. R. Brewer, M. Corzett, R. Balhorn, Protamine-induced condensation and decondensation of the same DNA molecule. Science 286, 120–123 (1999).

26. N. Makita et al., Salt has a biphasic effect on the higher-order structure of a DNA-protamine complex. J Phys Chem B 115, 4453–4459 (2011).

27. Y. Ke et al., 3D Chromatin Structures of Mature Gametes and Structural Reprogramming during Mammalian Embryogenesis. Cell 170, 367–381 e320 (2017).

28. M. Sun et al., Monitoring the compaction of single DNA molecules in Xenopus egg extract in real time. Proc Natl Acad Sci U S A 120, e2221309120 (2023).

29. X. Yan et al., Development of a mechanism-based, DNA staining protocol using SYTOX orange nucleic acid stain and DNA fragment sizing flow cytometry. Anal Biochem 286, 138–148 (2000).

30. D. Kota, R. Prasad, H.-X. Zhou, ATP Mediates Phase Separation of Disordered Basic Proteins by Bridging Intermolecular Interaction Networks. J Am Chem Soc, in press (2023).

31. C. G. Kalodimos et al., Structure and flexibility adaptation in nonspecific and specific protein-DNA complexes. Science 305, 386–389 (2004).

32. J. R. Horton, K. Liebert, S. Hattman, A. Jeltsch, X. Cheng, Transition from nonspecific to specific DNA interactions along the substrate-recognition pathway of dam methyltransferase. Cell 121, 349–361 (2005).

33. J. Iwahara, M. Zweckstetter, G. M. Clore, NMR structural and kinetic characterization of a homeodomain diffusing and hopping on nonspecific DNA. Proc Natl Acad Sci U S A 103, 15062–15067 (2006).

34. H. X. Zhou, Rapid search for specific sites on DNA through conformational switch of nonspecifically bound proteins. Proc Natl Acad Sci U S A 108, 8651–8656 (2011).

