## supplementary materials for "Multiple modes of DNA compaction by protamine"

Science
AAAS

### Supplementary Materials for

#### Multiple modes of DNA compaction by protamine

Vikhyaat Ahlawat[1] and Huan-Xiang Zhou[1]*

[1]Department of Chemistry and Department of Physics, University of Illinois Chicago, Chicago, United States.

**The PDF file includes:**

Materials and Methods
Figs. S1 to S8
References

**Materials and Methods**

Materials

Protamine sulfate (catalog # P4020) was from Sigma Aldrich. Sulfate was removed by dialysis (*30*) before use. A protamine variant, i.e., with FITC labeling, was also prepared (*30*). DRAQ5 (catalog # 62251) was from Thermo Scientific. SYTOX Orange, streptavidin-coated polystyrene beads (4.34 µm diameter), and phosphate buffered saline (PBS) as a kit (catalog # 00012) and biotinylated λ-DNA (catalog # 00001) were from LUMICKS. For confocal imaging, λ-DNA (catalog # N3011S) from New England Biolabs (NEB) was used. Imidazole buffer (10 mM, pH 7) was prepared by dissolving imidazole (catalog # 396745000, Thermo Scientific) in milli Q water.

Confocal microscopy

All experiments were conducted at room temperature on a LUMICKS C-Trap instrument, which combines a confocal fluorescence imaging module with dual-trap optical tweezers. For confocal imaging, a custom sample chamber was built with a clean glass coverslip attached to a microscope slide using double-sided tape. Each sample was 15 µL in volume, containing 2 ng/µL λ-DNA (NEB), 5 µM DRAQ5, and 0 to 10 µM protamine in imidazole buffer. For protamine-DNA colocalization imaging, protamine was doped with a FITC-labeled variant (9:1 ratio).

Confocal scanning was performed with 488 nm excitation for FITC and 638 nm excitation for DRAQ5 (8% laser power; 0.1 ms pixel dwell time; 0.1 µm pixel resolution). Images were exported in tiff format and processed using ImageJ. The maximum dimension ($D_{max}$) of DNA foci was measured as the diameter of a circumscribing circle.

We fit the dependence of $D_{max}$ on protamine concentration ([PM]) to a model where an increase in [PM] results in more coverage of the DNA molecule, and $D_{max}$ is the length of the still not covered portion. This model predicts

$$D_{max} = \frac{D^0_{max} - w_{point}}{1 + [PM]/K_{eff}} + w_{point} \qquad [S1]$$

where $D^0_{max}$ is the maximum dimension in the absence of protamine, $K_{eff}$ is an effective dissociation constant, and $w_{point}$ is the image size of a point source. This model is similar to the one used by Brewer et al. (*25*) to analyze single DNA molecule compaction kinetics, where the remaining DNA length at a given time is the length still not covered by protamine up to that time.

Single-molecule force spectroscopy

Force-extension curves were acquired using the dual-trap optical tweezers module, with sample components (600 µL each) loaded to microfluidics channels. The channels were filled with protamine (0.5, 1, or 10 µM in imidazole buffer), biotinylated λ-DNA (80 pg/µL in PBS), streptavidin-coated beads (0.004% w/v in PBS), and buffer (PBS), respectively. The trapping laser was set to 20% overall power and a 50:50 split between trap 1 and trap 2, resulting in a stiffness of 160 pN/µm for each trap.

All channels were opened at 1 bar pressure for 2 minutes to fill up the laminar flow cell with the sample components. In the laminar flow, two beads were trapped from the bead channel and then brought to the DNA channel to form a tether assembly. Experiments proceeded only on tethers that contained a single DNA molecule, which was verified by stretching in the buffer channel to produce an expected ~60 pN force plateau for strand separation. A stretch-relax cycle was then performed on the naked DNA to serve as reference, with the extension changing between 7.5 μm and ~17.0 μm (just before strand separation). Throughout the experiments, the speed of the moving trap (trap 1) was 1 μm/s.

The main protocol for stretch-relax cycles in the protamine channel started from a pre-stretch state (i.e., 7.5 μm extension, formed in the buffer channel and transferred to the protamine channel). The tethered DNA was stretched to ~17.0 μm (at the same maximum extension as in the reference), held in place for ~5 s (intra-cycle waiting period), and relaxed back to the pre-stretch state. Before the next cycle, the DNA was held in the pre-stretch state for ~20 s (inter-cycle waiting period), during which the protamine channel was opened under a mild flow (0.2 bar pressure). The stretch-relax cycles were repeated until the tether broke.

Variations of this protocol were introduced to prove different points. For example, for Fig. S2 right panels, the stretch-relax cycle was carried out in the buffer channel; for Fig. 4A-G and fig. S6, the stretch in the fourth cycle proceeded to ~20 μm to produce strand separation. For Fig. 4H, such an overstretch was first performed in the buffer channel. The DNA was then brought to the protamine channel and the flow was opened at 0.2 bar pressure for 20 s. Subsequently the DNA was relaxed to the pre-stretch state. After a waiting period of ~5 s, the DNA was finally stretched to ~17 μm. Note that the strand-separation force in the buffer channel is ~5 pN higher than in the protamine channel, due to the presence of ~150 mM salt in the PBS buffer; salt is known to increase the strand-separation force (*35*).

Correlative force-fluorescence measurements

These experiments were similar to those in the preceding subsection, except that SYTOX Orange was included. In the main protocol, this dye (0.5 μM) was premixed into the protamine (10 μM) channel. Kymographs or 2-D scans were acquired simultaneously with force-extension curves (Fig. 3). For fig. S7C (also presented as fig. S8D) and fig. S8B, the dye was in a separate channel as opposed to premixed into the protamine channel. For fig. S7B, D, 2D scans and kymographs were acquired after the DNA was transferred to the buffer channel. Imaging was performed with 532 nm excitation for SYTOX Orange (30% laser power; 0.1 ms pixel dwell time; 0.1 μm pixel resolution). Force data were processed using Matlab; images were processed using ImageJ.

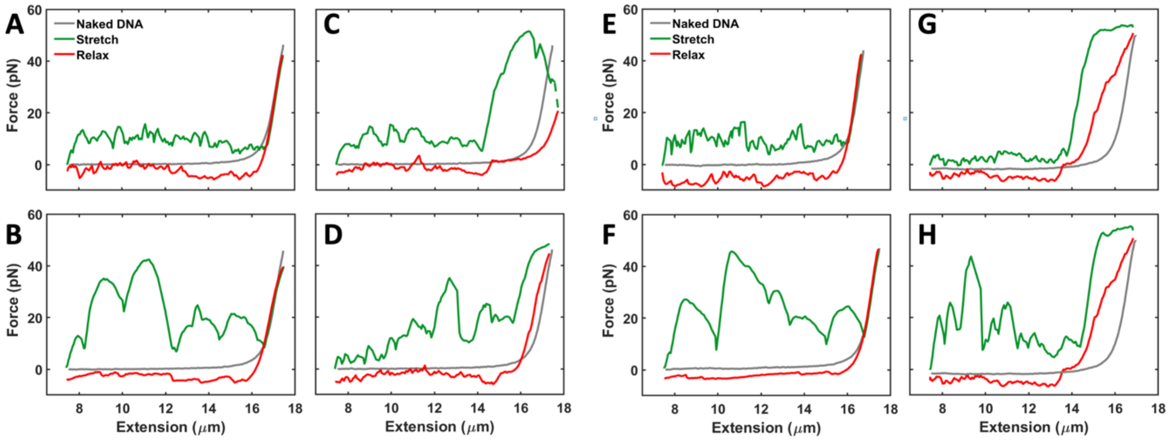


**Fig. S1. Force-extension curves demonstrating different modes of λ-DNA compaction by protamine.** (**A**-**D**) Same as Fig. 2A-D, but protamine is at 0.5 μM. (**E**-**H**) Same as Fig. 2A-D, but protamine is at 1 μM.

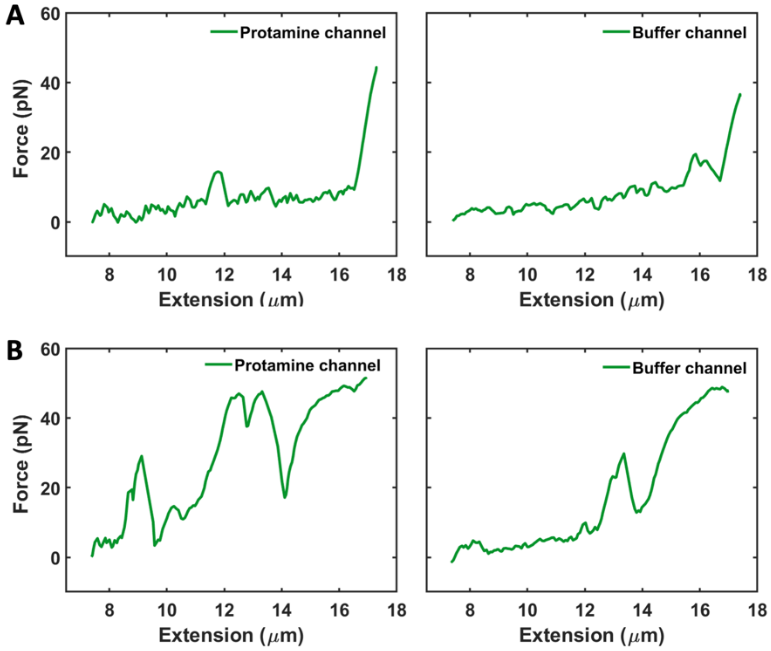


**Fig. S2. Protamine remains bound when DNA is transferred to the buffer channel.** (**A**) A protamine-bound DNA molecule showing the bending mode continues to show the same mode after transferring to the buffer channel. (**B**) After forming protamine-mediated coils, a DNA molecule repeats the coiling mode in the buffer channel.

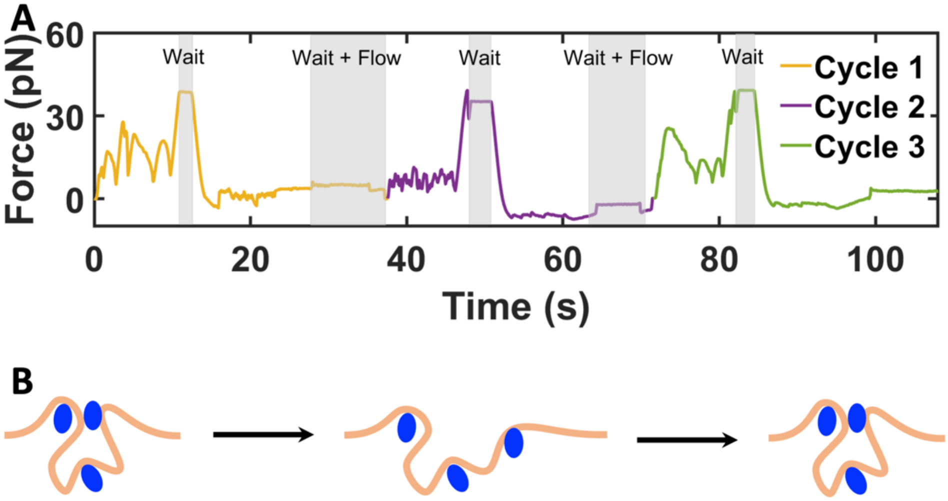


**Fig. S3. Interconversion between the bending mode and looping mode.** (**A**) Pulling force in three successive stretch-relax cycles. The continuous rise in pulling force at high stretch (just before the intra-cycle waiting period) indicates the compaction mode is either bending or looping; these two modes can be distinguished by the maximum forces at low stretch: near 30 pN for the looping mode (cycles 1 and 3) but near 10 pN for the bending mode (cycle 2). In the inter-cycle waiting period, protamine flow was open. (**B**) Illustration of the conversion between the looping and bending modes. DNA and protamine are represented by an orange line and blue ovals, respectively.

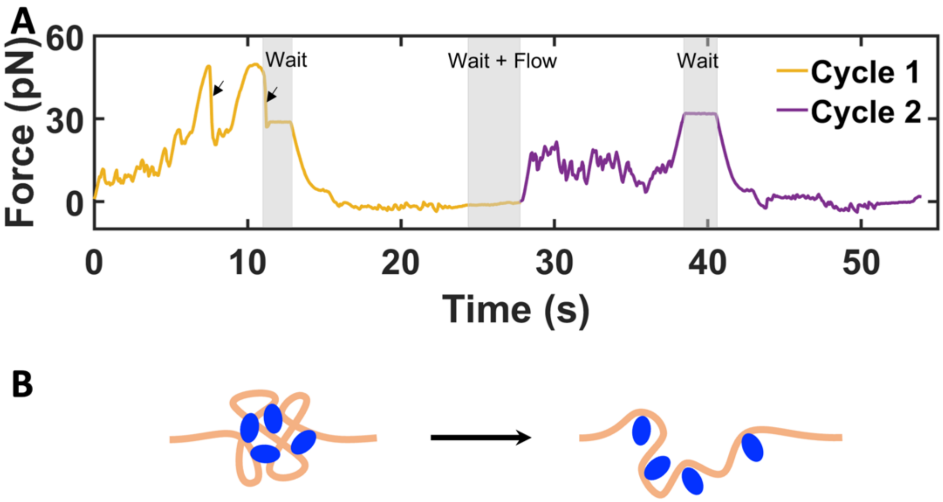


**Fig. S4. A rare conversion from the coiling mode to the bending mode.** (**A**) Pulling force in two successive stretch-relax cycles. A plateau around 50 pN at high stretch indicates the coiling mode in cycle 1, whereas a continuous rise at high stretch, along with a maximum force of ~20 pN at low stretch, indicates the bending mode in cycle 2. Arrowheads in cycle 1 highlight large drops in force in the plateau region and in the subsequent waiting period, signifying the unraveling of coils that leads to the bending mode in the next cycle. (**B**) Illustration of the conversion from the coiling mode to the bending mode.

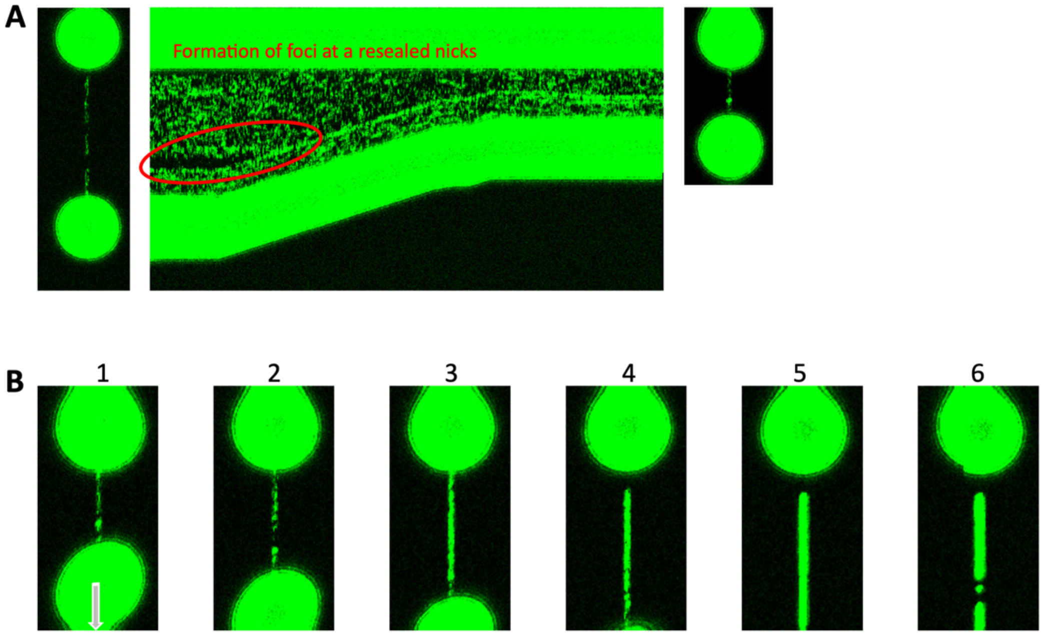


**Fig. S5. Conversion from the bending mode to the coiling mode.** (**A**) A kymograph showing that, after the stretch of a DNA molecule in the bending mode, nicks develop in the subsequent waiting period but reanneal as the DNA is relaxed. As highlighted by a red oval, at the reannealed locations, SYTOX Orange foci emerge and then persist as the DNA is relaxed back to the pre-stretch state. A 2D scan just before the kymograph also captures the nicks whereas a 2D scan right after the kymograph captures the SYTOX Orange foci. (**B**) A series of 2D scans capturing the coiling mode in the next stretch. Note that the images of the bead at the bottom are distorted because it moved at a speed of 1 μm/s while the scanning took place.

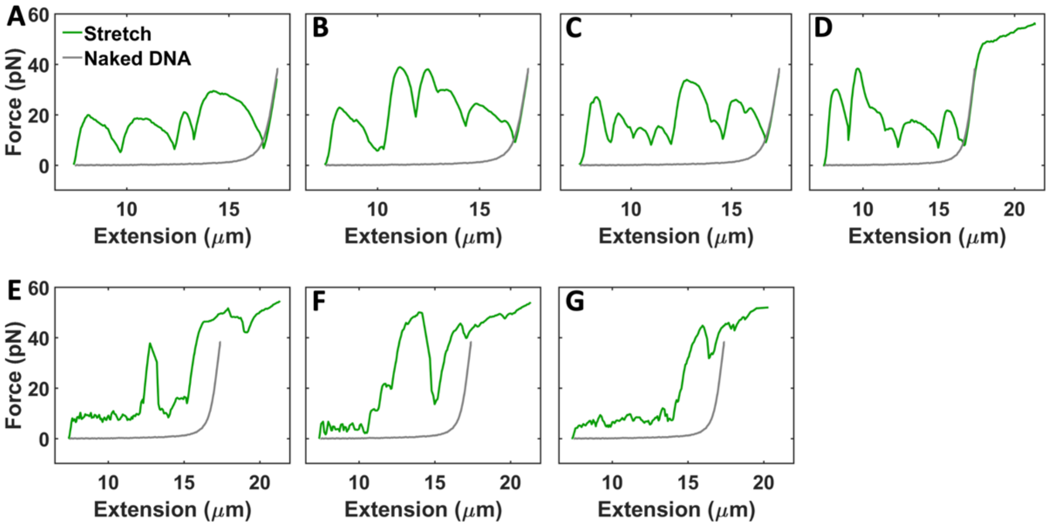


**Fig. S6. Induction of the coiling mode by overstretching DNA.** (**A-G**) A seven-cycle sequence for inducing the conversion from the looping mode to the coiling mode. (A-C) present three successive regular stretch-relax cycles in the looping mode; overstretch is introduced in (D), resulting in the coiling mode in the subsequent cycles shown in (E-G).

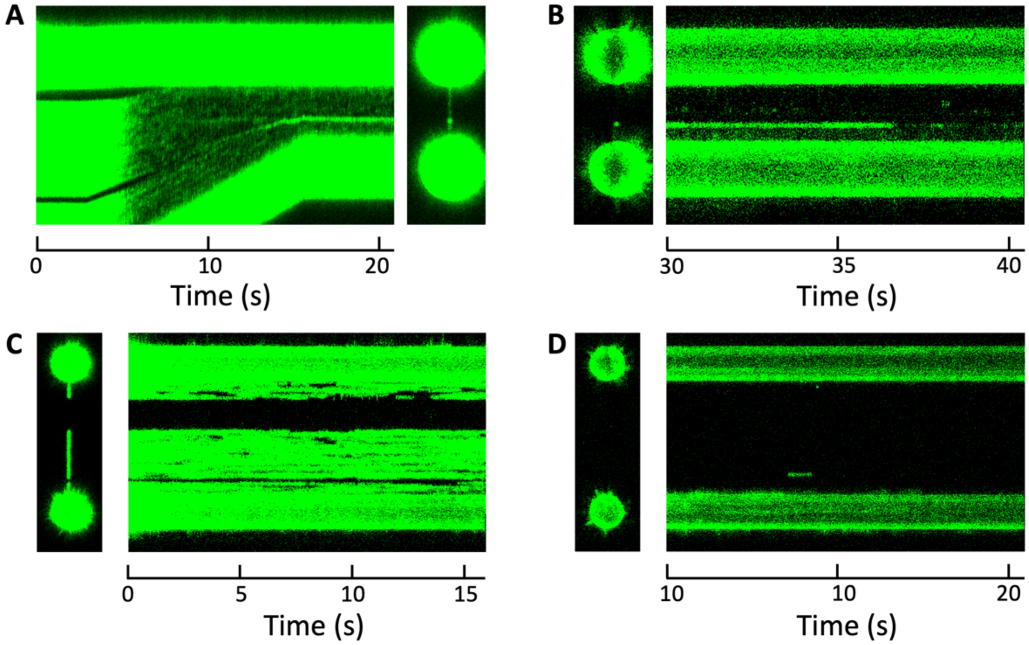


**Fig. S7. Trapping of intercalated SYTOX Orange by protamine in coils.** (**A**) Relax of a coil-containing DNA molecule back to the pre-stretch state, where SYTOX Orange foci are visible. Note that this coiling mode was produced by overstretching in the buffer channel (protocol of Fig. 4H). (**B**) After transferring the DNA molecule to the buffer channel, SYTOX Orange foci persist for > 35 s and then fade. Note that the kymograph starts at 30 s post-transfer. (**C**) As a negative control, a naked DNA molecule is overstretched and then stained with SYTOX Orange. (**D**) As soon as the stained DNA is transferred to the buffer channel, the dye is completely released. The 2D scan is right after the transfer, followed by the kymograph starting at 10 s post-transfer.

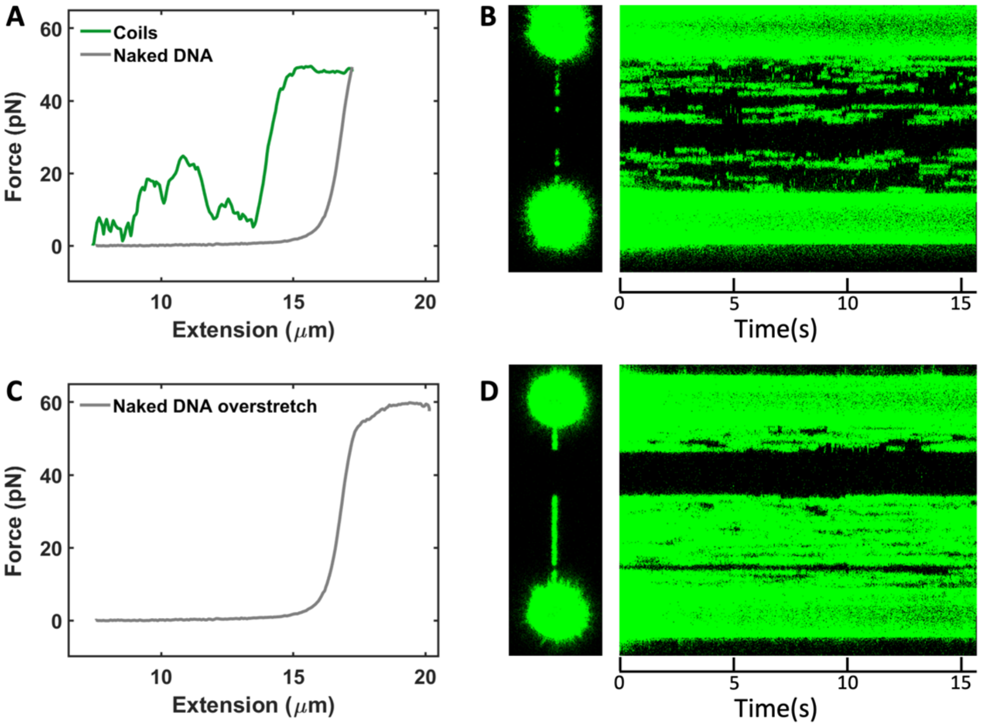


**Fig. S8. Resistance of SYTOX Orange intercalation by protamine pre-bound in coils.** (**A**) Stretch of a DNA molecule exhibiting the coiling mode. (**B**) The overstretched, coil-containing DNA molecule is transferred to the SYTOX Orange channel, where dye staining occurs only intermittently. (**C**) As a positive control, a naked DNA is overstretched. (**D**) When the overstretched naked DNA molecule is transferred to the SYTOX Orange channel, staining occurs continuously and uniformly in unnicked regions. Panel (D) is the same as fig. S7C.

**Supplementary References**

35. J. R. Wenner, M. C. Williams, I. Rouzina, V. A. Bloomfield, Salt dependence of the elasticity and overstretching transition of single DNA molecules. *Biophys J* **82**, 3160-3169 (2002).
